# CytoGAN: Generative Modeling of Cell Images

**DOI:** 10.1101/227645

**Authors:** Peter Goldsborough, Nick Pawlowski, Juan C Caicedo, Shantanu Singh, Anne E Carpenter

**Author notes:** This work has been conducted as part of an internship at the Imaging Platform of the Broad Institute. 31st Conference on Neural Information Processing Systems (NIPS 2017), Long Beach, CA, USA.

## Abstract

We explore the application of Generative Adversarial Networks to the domain of morphological profiling of human cultured cells imaged by fluorescence microscopy. When evaluated for their ability to group cell images responding to treatment by chemicals of known classes, we find that adversarially learned representations are superior to autoencoder-based approaches. While currently inferior to classical computer vision and transfer learning, the adversarial framework enables useful visualization of the variation of cellular images due to their generative capabilities.

## 1 Introduction

Advances in high-throughput microscopy systems have enabled acquisition of large volumes of high-resolution cell images [3]. This paves the way for novel computational methods that can leverage these large quantities of data to study biological systems. Our work focuses on the task of morphological profiling, which aims to map microscopy images of cells to representation vectors that capture salient features and axes of variation in an unsupervised manner [4]. These representations ideally divide the morphological space into clusters of cells with similar properties or, in the case of chemical or genetic treatments added to the cells, similar function.

Current techniques for morphological profiling broadly fall in two categories: a) classical image processing, using specialized software like CellProfiler [5] to capture representations via manually-tuned segmentation and traditional computer vision pipelines, and b) transfer learning to extract features learned by deep convolutional neural networks originally trained to classify miscellaneous objects [16, 1]. Classical computer vision approaches offer better interpretability of features but require more human tuning of the segmentation algorithms, and are limited by the feature set implemented in the image analysis software. Current transfer learning techniques have been shown to outperform classical methods in at least one dataset [16, 1]. However, given that these networks were trained on natural images (in RGB), they do not discover the relations of the biologically meaningful image channels. Instead, their superior performance is likely due to their ability to extract high-level vision features, which appear to capture the overall structure of cells, but not necessarily all the intricate details of their morphological variations. Therefore, we hypothesized that learning representations specifically adapted to cell images would be valuable.

We employ Generative Adversarial Networks (GANs) [7] to build a generative model of single cell images from the BBBC021 dataset [12]. We show that this model learns rich feature representations and synthesizes realistic images that are useful for exploring morphological variation in cells. The main advantages of this method are:

- *Adaptability:* Because this method learns from its training data it is able to disentangle factors of variation and extract inherent semantic relationships. Transfer learning approaches lack this ability and cannot capture the intrinsic relations between the biologically meaningful channels.
- *Translating into biological phenotypes:* The generative abilities of this method enable useful visualizations of cells to help translate data variations into biological phenotypes.

## 2 Related Work

Our work lies at the intersection of automated morphological profiling, and representation learning with deep neural networks, in particular generative architectures. Caicedo et al. [3] recently outlined the state and challenges of the morphological profiling problem. Prior to this, Ljosa et al. [11] compared the performance of various dimensionality reduction techniques for CellProfiler features on the BBBC021 benchmark [12], consisting of MCF7 cells exposed to different chemical treatments. Pawlowski et al. [16] for the first time reported a representation-learning method based on deep learning that is competitive with hand-engineered features at the task of predicting mechanism-of-action (MOA) of chemicals. Ando et al. [1] also applied transfer learning with a different architecture and introduced a novel feature normalization method. Other related work on this dataset include supervised classification [10], transfer learning on CellProfiler features [9], and dimensionality reduction using autoencoders [19].

Few published studies have applied unsupervised deep learning techniques to the task of feature extraction in morphological profiling. Pawlowski [15] first investigated autoencoder-based methods but reported results far inferior to hand-tuned features or transfer learning approaches. Our model is more related to the work by Osokin et al. [14] and Johnson et al. [8], wherein GANs were used to model cell images, although their applications did not include morphological profiling.

## 3 Using GANs for Representation Learning

Goodfellow et al. [7] introduced GANs as a game of two players: a *generator G* and a *discriminator* or *critic D.* The former receives samples **z** drawn from a noise prior *P*_noise_ which it maps to values *G*(**z**) that should resemble elements of some data distribution *P*_data_. The discriminator must learn to distinguish such synthetic samples from real values x ~ *P*_data_. The critic’s confidence in the realism of the generator’s productions is used as feedback to *G*, guiding it to synthesize ever more realistic replicates of samples from the data prior. This procedure is formalized in a zero-sum game,

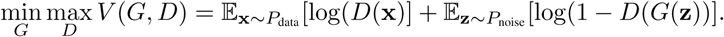

Radford et al. [17] first specialized GANs to image synthesis by introducing Deep Convolutional GANs (DCGANs). DCGANs implement the generator and discriminator based on convolutional operations. Derivations of DCGANs such as Least Squares GAN (LSGAN) [13] or Wasserstein GAN (WGAN) [2] tackle instabilities in the training procedure of early generative adversarial models. In our experiments, we found LSGAN to be most stable, in part leading to higher quality generated images than both DCGAN and WGAN.

The original GAN framework does not include an explicit means of performing inference. As such, we require extensions that allow mapping of a sample x drawn from the data prior to a latent representation via some encoder transformation *E*(x). A common approach is to interpret the penultimate layer of the discriminator as a latent space. The activations of this layer serve as a source of representation vectors. These latent codes are thought to be meaningful because the discriminator must develop a strong internal representation of its input to succeed at its discrimination task. Furthermore, this method imposes no computational overhead compared to vanilla GANs. Figure 1 shows a simplified schematic of our GAN architecture. More advanced encoders could tie this encoding to a corresponding noise vector.

**Figure 1:**
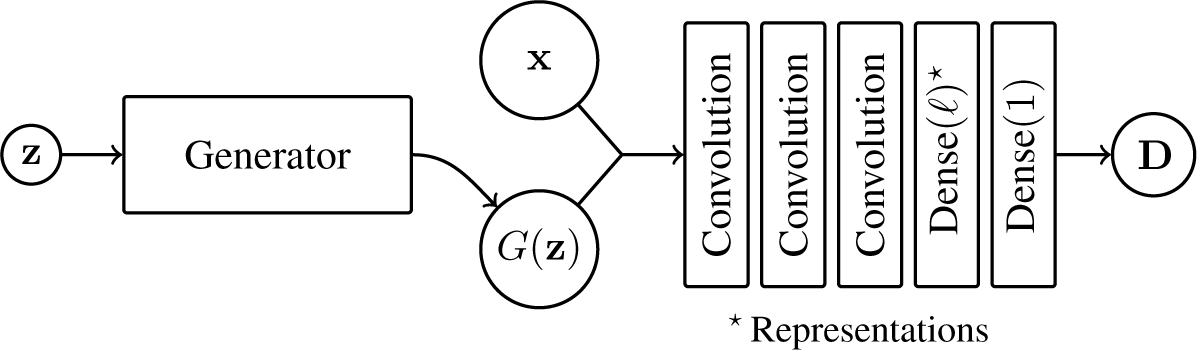
A simplified diagram of the modified DCGAN [17] architecture we employ for our experiments. On a high level, the discriminator is a sequence of convolution layers, leading to a fully connected (dense) layer with ℓ neurons. We interpret the activations of these ℓ neurons as latent vectors and further feed them into a final dense layer with just one activation, forming the discriminator’s output.

## 4 Exploring Biological Phenotypes Using Cell Image Synthesis

GANs have been proven to synthesize realistic images both within and beyond the biological domain [7, 17, 14]. We examine if this ability transfers to cell images extracted from the BBBC021 dataset of human breast cancer cell lines. Each image in BBBC021 consists of three *channels* corresponding to DNA, F-Actin and *β*-Tubulin, cellular components visualized by fluorescence microscopy. We stack these channels and treat them as RGB images via a simple DNA ↦ R, *β*-Tubulin ↦ G, F-Actin ↦ B mapping. We obtain 1.3 million single cell images by segmenting the raw images using CellProfiler. Finally, we normalize the luminance of each channel to the range [0 − 1].

Figure 2 shows examples of images generated with LSGAN, WGAN and DCGAN architectures alongside real images. The synthetic images, particularly those produced by LSGAN, are not only highly detailed and realistic, but also consistent with their biological nature. For example, it is characteristic that *β*-Tubulin (green) forms a circular halo cradling the nucleus. This property is maintained clearly in most generated images.

**Figure 2:**
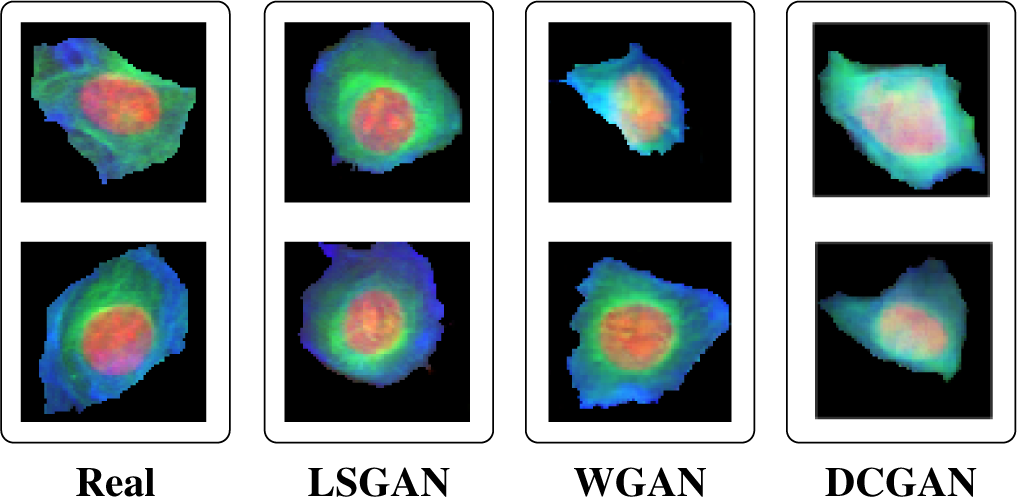
Examples of real BBBC021 cell images juxtaposed with synthetic cell images generated with LSGAN, WGAN and DCGAN.

To assess the quality of synthesized images we presented three expert biologists with 50 real cell images and 50 artificial cells generated with LSGAN. We randomized the order of images and asked the experts to judge whether each cell was real or fake. On average, 30% of the time the synthetic cells were realistic enough to fool the human jurors into believing they were real.

Current approaches to morphological profiling extract features that are challenging to translate into biological meaning, such as Zernike moments. While other features capture readily understandable concepts such as the area occupied by the nucleus, even these are difficult to interpret when a given class of cells is defined by several such features in combination. For transfer learning it is nearly impossible to visualize the concept of a feature. In contrast, the noise space of GANs is known to be highly interpretable and reveal rich semantic structure [17]. We are able to demonstrate this for images of cells. Figure 3 exhibits how interpolating between two noise samples leads to smooth transitions in synthesized cells. This supports the fact that the generator learned a low-dimensional manifold of the images. Figure 4 shows that the noise space encodes semantic relationships, enabling algebra on interpretable properties of the images such as size of the nucleus or *β*-Tubulin content. While there is no way to encode images into *P*_noise_ with the methods presented so far, we believe more advanced architectures that enable this will be highly valuable if they can maintain these remarkable semantic properties.

**Figure 3:**
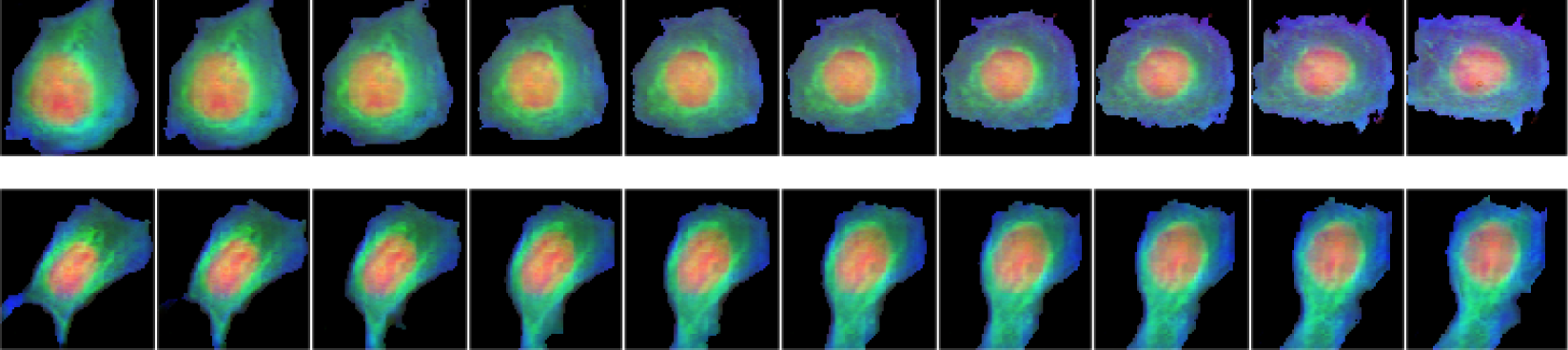
Interpolating between two points **z**_1_, **z**_2_ ~ *P*_noise_ results in smooth transitions in the synthesized cell images. Each row shows the transition from **z**_1_ to **z**_2_ from left to right.

**Figure 4:**
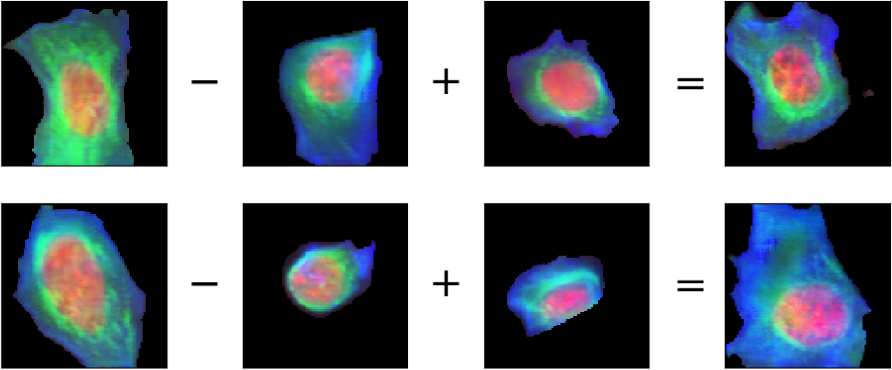
Vector algebra in the noise space translates to biologically valuable relationships in generated images. In the first row, subtracting the vector for a cell with small quantities of *β*-Tubulin (green) from one with high amounts yields a vector representation for “higher *β*-Tubulin content”. In the second row, the difference between a large and small nucleus encodes the semantic meaning of “larger nucleus”, which can be added to vectors of other cell images to grow the size of the nucleus.

## 5 Representation Learning for Morphological Profiling

We test the quality of representations extracted via the discriminator by evaluating their ability to cluster treatments of similar function, here the mechanism-of-action. We obtain treatment profiles by averaging the extracted single cell representations found as intermediate activations of the discriminator. Further, we follow the experimental protocol of [11] and report an average MOA classification accuracy of a leave-one-compound-out cross validation using a one nearest neighbor classifier.

Table 1 compares the accuracy of our technique to classical CellProfiler features and transfer learning. We found that the quality of representations extracted by our approach is not yet competitive with other methods. Nevertheless, we believe that the adversarial approach has significant benefits regarding its ability to translate into biological phenotypes as outlined above. Additionally, this framework is able to adapt to the dataset at hand and extract inherent relations that are not captured by previous techniques. We believe that further improvements are possible, as the quality of synthesized images correlates with classification accuracy. Accordingly, the best performing method (LSGAN) also yields the highest image quality, as judged qualitatively by expert biologists and shown in Figure 2.

**Table 1:** Comparison of the mechanism-of-action classification accuracy. CP refers to the results by Singh et al. [18] using CellProfiler features. The Deep Transfer methods correspond to Deep Feature Transfer as proposed by Pawlowski et al. [16] and refined by Ando et al. [1].

| DCGAN | WGAN | LSGAN | VAE [15] | CP [18] | Deep Transfer [16] | Deep Transfer [1] |
| --- | --- | --- | --- | --- | --- | --- |
| 51 % | 56 % | 68 % | 49 % | 90 % | 91 % | 96 % |

## 6 Conclusion

This work investigates the use of GANs for the domain of cell microscopy imaging, in particular morphological profiling. First, we demonstrate the abilities of our generative model with the exploration of morphological phenotypes by synthesizing realistic images of cells and performing transformations on them such as interpolation and vector algebra. Second, we extend standard GANs with an encoder and assess the quality of learned representations by evaluating their mechanism-of-action classification performance following [11]. Even though adversarially learned representations are currently inferior at this task, we argue that further enhancements to GANs and encoding schemes will lead to biologically richer latent representations and better MOA classification accuracy.

This work covers only a small fraction of possible applications of GANs to the domain of microscopy images. For example, we hope that future work investigates BiGANs[6] or other novel solutions to infer latent features with GANs. We believe that improved inference, combined with the interpretable nature of the GAN framework, may enable simulated experiments by performing algebra with vectors corresponding to cell lines, diseases or other perturbations.

## Acknowledgements

We thank Allen Goodman, Jane Hung, Claire McQuin, Kyle Karhohs, Beth Cimini, Tim Becker, Minh Doan, Jeanelle Ackerman and Ray Jones of the Broad Institute for fruitful discussion, guidance and support with our experimental setup. We also extend our gratitude to Mike Ando and Google for providing computational resources to accelerate our research.

This work was supported in part by a grant from the US National Science Foundation (CAREER DBI 1148823 to AEC).

Nick Pawlowski is supported by Microsoft Research through its PhD Scholarship Program and the EPSRC Centre for Doctoral Training in High Performance Embedded and Distributed Systems (HiPEDS, Grant Reference EP/L016796/1).

